# Optimizing MR-based gaze-decoding for eyes-closed eye-tracking in fMRI

**DOI:** 10.64898/2026.07.07.736972

**Authors:** Sina M. Kling, Uriel Lascombes, Matthias Nau, Guillaume S. Masson, Martin Szinte

## Abstract

Eye movements provide valuable insights into human cognition and are a critical variable in numerous functional magnetic resonance imaging (fMRI) studies. Yet, when the eyes are closed, camera-based eye-tracking is unavailable, making studies of eyes-closed states challenging. Here, we address this gap using DeepMReye, a deep learning framework for camera-free gaze reconstruction from the MR-signal of the eyes. We first show that fine-tuning DeepMReye on visuomotor calibration data acquired with the eyes open significantly improves gaze decoding, and that this fine-tuning does not require simultaneous camera-based data. We next assessed whether decoding could be extended to eyes-closed states using a novel auditory-guided task, in which participants gazed at learned target positions with and without visual input, and with their eyes open, blinking, or closed. While DeepMReye was originally trained exclusively on eyes-open data, the network successfully generalized to eyes-closed periods, and this generalization was further improved through task-specific fine-tuning. Finally, fine-tuning also improved decoding of eyelid-state (open versus closed) directly from the MR-signal. These findings demonstrate that gaze and eyelid-state monitoring during eyes-closed periods is feasible, enabling broader integration of eye-tracking in fMRI research.

## Introduction

Eye movements offer a dynamic window into perceptual and cognitive processes. As essential components of visual behavior, they not only indicate where we direct our gaze, but also reflect how we allocate attention, process information, and make decisions (McCarley & Kramer, 2007). Accordingly, eye movements have been linked to language comprehension and memory (Hannula et al., 2010; Mézière et al., 2023; Ryan & Shen, 2020), attention and decision-making (Blair et al., n.d.; Fiedler & Glöckner, 2012; Gidlöf et al., 2013; Tatler, 2007), and mental imagery (Benedek et al., 2017; Laeng et al., 2014). Eye movements are informative well beyond visual processing, extending even to sleep, where rapid eye movements during REM reflect internal cognitive activity in the absence of any external stimulus (Senzai & Scanziani, 2022).

In functional magnetic resonance imaging (fMRI) research, eye movement monitoring is essential. Tracking gaze ensures participants attend to stimuli as intended, helping to reduce confounds from unintended viewing behavior. Even subtle gaze shifts can substantially alter activation patterns in visual cortex, particularly in paradigms relying on retinotopic mapping or precise stimulus presentation (Laconte et al., 2007; Lee et al., 2015). Access to accurate gaze data is thus vital for controlling attention, verifying task compliance, and improving the interpretability of fMRI findings. Traditional MR-compatible eye-tracking systems typically rely on infrared video to track corneal reflections and pupil position. However, in addition to their high cost, these camera-based systems face practical limitations in MRI environments, including constrained setup geometry, interference from scanner hardware, signal dropout, and susceptibility to motion artifacts. Most critically, they require participants’ eyes to remain open, making them unusable in contexts where eyes are closed or obscured such as during sleep, eyes-closed mind wandering and other resting state paradigms. To overcome these constraints, recent efforts have explored decoding gaze directly from eye-voxel signals. Early approaches demonstrated that eye movements could be inferred from the MR signal of the eyes itself (Beauchamp, 2003; Tregellas et al., 2002), with subsequent work extending to gaze decoding under naturalistic viewing conditions (Nau et al., 2025; Son et al., 2020). DeepMReye, an established framework for eye voxel-based gaze decoding, uses convolutional neural networks to reconstruct gaze from MRI signals of the eyeballs and surrounding tissues (Frey et al., 2021). This camera-less approach performs robustly across diverse viewing tasks and has the unique ability to be applied retrospectively to datasets lacking camera-based eye-tracking data.

However, DeepMReye has been validated mostly under eyes-open conditions, with only limited evidence of successful performance under eyes-closed conditions (Frey et al., 2021). However, many fMRI paradigms include periods when participants’ eyes are closed, such as resting-state scans, sleep studies, visual-imagery tasks, and clinical protocols (Bianciardi et al., 2009; Fulford et al., 2018). This creates a significant gap, as eye movement dynamics may differ substantially between eyes-open and eyes-closed states (Allik et al., 1981; Collewijn et al., 1985), and eyelids closure can influence eyeball position and motion (Kirchner et al., 2022). The present study addresses this gap with two primary aims. First, we determined how DeepMReye can be further optimized for specific experimental set-ups, as our local one, by systematically fine-tuning the network to local scanner environments and task-specific viewing dynamics. Second, we develop a novel paradigm using auditory-guided eye movements to fine-tune the model for usage without visual input and with eyes blinking or closed, further called visual/eyes state tasks. Our results demonstrate that fine-tuning with task-specific eyes-open data significantly improves decoding performance of DeepMReye during “guided fixation”, “guided pursuit” and “free viewing” parts. Furthermore, we demonstrate that task-specific fine-tuning substantially improves classification accuracy during eyes-closed states. Finally, we show that this fine-tuning also improves decoding of eyelid-state directly from the MR-signal. This advancement enables gaze monitoring in previously inaccessible fMRI contexts and opens new possibilities for retrospective analysis of large existing fMRI data resources. By extending MR-based gaze decoding beyond its current limitations, this work provides researchers with a powerful tool for understanding eye movements across the full spectrum of conscious states.

## Material and methods

### Participants

Seventeen participants (9 females, 8 males) took part in the experiment (ages 24–39). All except the authors (N=3, S.M.K, U.L, M.S) were naïve to the purpose of the study, and all had normal or corrected-to-normal vision. Two participants were excluded after the training phase due to poor task performance or their decision to withdraw because of task difficulty. The remaining 15 participants completed all runs of both the visuomotor calibration and visual/eyes state tasks. Participants were compensated for their participation in the study.

### MRI data acquisition

T1-weighted (0.8 mm isotropic resolution) and T2-weighted (0.8 mm isotropic resolution) structural images were acquired for each participant using MPRAGE at the MRI center of the Institute de Neurosciences de la Timone on a Siemens Prisma 3T scanner (Siemens Healthineers, Erlangen, Germany) with a 64-channel receive coil array. Functional images were collected using a 2D gradient-echo EPI sequence with a multiband acceleration factor of 4, at 2 mm isotropic resolution, 1.2 s TR, and 60 axial slices covering the entire brain. The field of view was carefully adjusted to fully include the eyeballs, ensuring complete sampling of ocular structures. To correct for susceptibility-induced distortions, particularly those that can alter the apparent shape and signal profile of the eyeballs, we acquired spin-echo EPI field maps in both anterior-to-posterior (AP) and posterior-to-anterior (PA) phase-encoding directions. These dual-polarity fieldmaps enabled accurate estimation of distortion fields using *topup* algorithm and were critical for ensuring precise reconstruction of the MR signal in and around the eyes.

### Experimental sessions, apparatus and tasks

The experiment consisted of two sessions held on separate days. In the first session, participants were trained on the tasks outside the scanner. The second session took place in the MRI scanner and comprised two consecutive sets of tasks. Visuomotor calibration tasks (3 functional runs of ∼3 min each) and visual/eyes state tasks (3 functional runs of ∼6 min each). Together with preparatory scans, field inhomogeneity mapping, and structural image acquisition, a scanner session took approximately 1 hour in total.

Stimuli were presented at a viewing distance of 120 cm, on a projection screen situated at the end of the bore (20 dva horizontally by 20 dva vertically) and viewed through a mirror placed on the head receiver. The screen had a spatial resolution of 1080 by 1080 pixels and a refresh rate of 120 Hz, back-projected with a PROPixx projector (VPixx Technologies Inc., Saint Bruno, Quebec, Canada) located outside the scanning room. The experimental software controlling the display, the sounds and the response collection was implemented in Matlab (The MathWorks, Natick, MA, USA) using the Psychophysics Toolbox (Brainard, 1997; Pelli, 1997). Participants gaze position was recorded using an infrared-video based eye-tracker (Eyelink 1000 system, SR Research, Ottawa, Canada). A 13-point custom calibration of the eye-tracker was performed before each task, and across runs if needed. Auditory stimuli were delivered using the OptoActive Active Noise Cancelling Headset System (OptoAcoustics Ltd., Mazor, Israël), which combines passive attenuation through acoustic padding with real-time algorithmic active noise cancellation. The active noise cancellation is synchronized to the MRI scanner allowing optimal attenuation during functional MRI echo-planar imaging sequences.

The visuomotor calibration tasks, comprised classical oculomotor parts, visually guided fixations, smooth pursuit eye movements, and free viewing of natural images, chosen because they drive robust, well-calibrated eye movements similar to those used in the initial training of DeepMReye. However, these tasks are limited to eyes-open data.

The visual/eyes state tasks, was therefore designed to extend this to structured eyes-closed eye movements. To both facilitate learning for participants and to systematically sample the space of possible visual and eye states, this part comprised four parts crossing two factors: the presence or absence of visual input, and whether participants’ eyes were open or closed.

### Visuomotor calibration tasks

The visuomotor calibration tasks were composed of three consecutive parts: “guided fixation”, “guided pursuit”, and “free viewing” (Figure 1A). These parts were preceded and followed by 6.0 s intervals of simple fixation in which participants were instructed to fixate a white bull’s eye (0.25 dva radius) on a black background shown at the screen center. The “guided fixation” part was composed of 50 trials of 1.2 s each, during which participants were instructed to fixate the bull’s eye presented at 25 positions (5 rows by 5 columns) forming a virtual square of 18 dva-side. Each of the 25 positions were repeated twice per run, they were never played in a fixed sequence and started and ended at the screen center. The “guided pursuit” part was composed of 54 trials of 1.2 s each during which the bull’s eye smoothly moved along a linear trajectory varying in amplitude and rotation angle across trials. The amplitudes were randomly chosen between 3, 5 and 7 dva. The angles were randomly chosen between 0 and 340 degrees of rotation (in steps of 20 degrees). The sequence of trials was selected such that the bull’s eye started and finished in the screen center and remained within a virtual square boundary of 18 dva-side. The “free viewing” part was composed of 10 trials of 3.6 s each during which participants visually explore 10 natural images (9 dva-square) presented in a random order. The images were randomly selected from a publicly available dataset (https://unsplash.com/). The same set of images were presented in random order across runs and participants.

**Figure 1.**
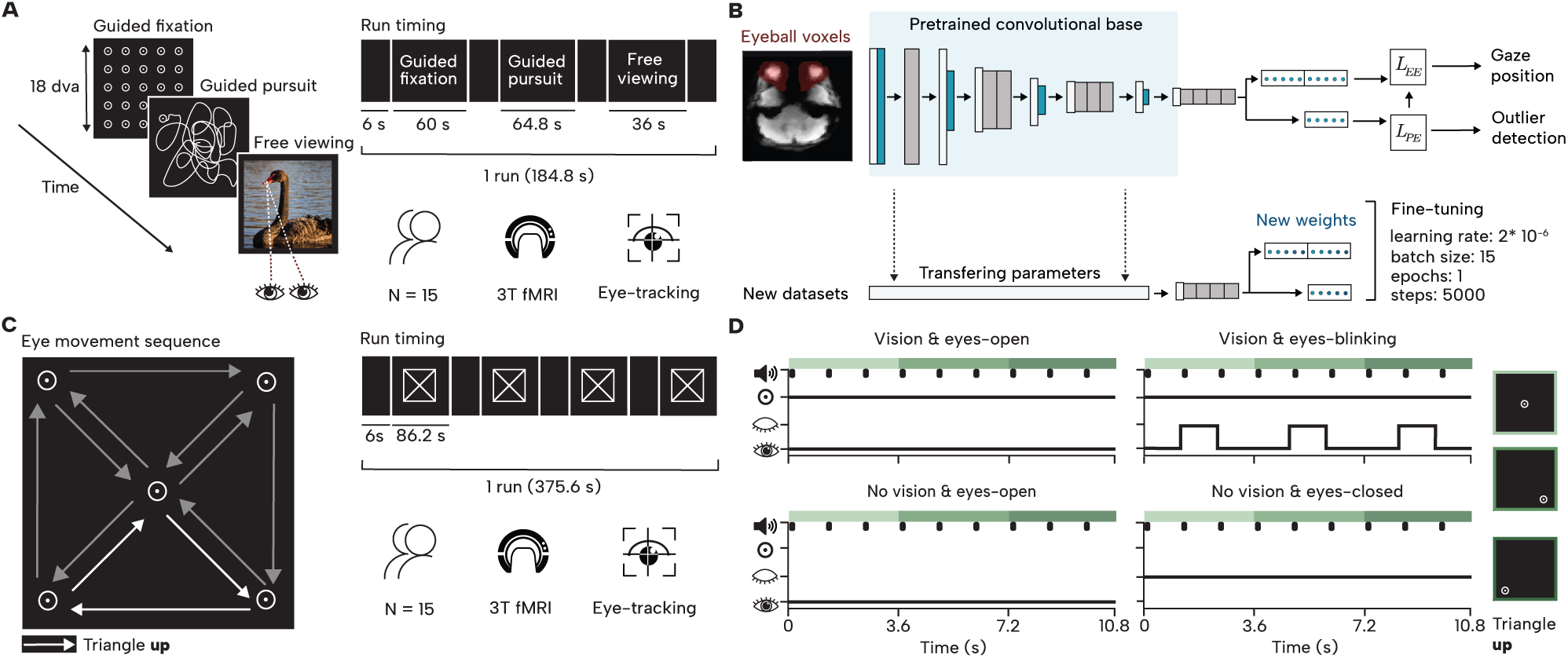
**A**. Visuomotor calibration tasks. Participants completed three consecutive parts (see Material and methods): guided fixation, guided pursuit, and free viewing of natural images. **B.** DeepMReye CNN architecture and fine-tuning procedure. DeepMReye takes eye mask voxels from BOLD images as input and predicts 2D gaze position through a series of convolutional layers, trained using two loss functions: Euclidean Error (EE) and the discrepancy between EE and a predicted error (PE). Fine-tuning was performed by retraining the network on new participant-specific input data and gaze labels. **C.** Visual/eyes state tasks. Participants fixated on a bull’s eye target across five consecutive screen positions (central and four corners of a virtual square) forming four triangle patterns with their gaze trajectories. **D.** The eye movement sequence was completed over four parts varying in visual input (vision/no vision) and eye state (open, blinked, closed), in which they followed a bull’s eye target guided by auditory tone sequences. Here is shown a single triangle sequence, forming a triangle pointing upwards.

### Visual/eyes state tasks

These tasks were composed of four consecutive parts manipulating both availability of visual information and eyes– or eyelid-state: “vision and eyes-open”, “vision and eyes-blink”, “no vision and eyes-open”, “no vision and eyes-closed”. These parts were preceded and followed by 6.0 s fixation intervals in which participants were instructed to fixate the bull’s eye on a black background shown at the screen center.

#### Vision and eyes-open

Participants performed a guided fixation task, with the eyes open, where they followed a bull’s eye target that appeared at five positions: the center of the screen and the four corners of an 18 dva virtual square centered on the screen (Figure 1C). Starting from the center position, participants tracked the bull’s eye as it moved to different positions every 3.6 seconds, tracing out triangular paths by connecting the center with pairs of corner positions. The bull’s eye movement sequence created four triangular patterns in order: an upward triangle, followed by a rightward triangle, a downward, and finally a leftward triangle. This entire sequence was performed twice, resulting in participants following eight triangular patterns in total. Trial timing (Figure 1D) was structured using auditory cues: a sequence of three ascending tones (440 Hz, 660 Hz, and 880 Hz, each 0.3 s in duration and separated by 900 ms intervals) signaled trial events, while the task began and ended with a 6.0 s inter-trial interval marked by five 300 Hz tones (0.3 s and separated by 0.9 s interval). By doing this, participants learned sequences of eye positions forming 4 triangles, that can then be used as memory/imaginary targets for the other tasks parts.

#### Vision and eyes-blink

Participants fixated on a bull’s-eye target with its central dot removed. The trial sequence and timing were identical to those in the “Vision and eyes-open*”* part, except that participants were instructed to close their eyes on every trial, performing a prolonged blink, between the second and third tone. This part served two purposes: to facilitate participants’ learning of the fixation sequence by providing an intermediate step between fully guided viewing and the subsequent parts in which visual input was absent, and to introduce natural, prolonged blink behavior into the dataset.

#### No vision and eyes-open

Participants maintained fixation at the same positions as in the previous parts, but no visual stimulus was presented. The auditory cues and timing structure were identical. Participants were instructed to use the tones to guide their fixation timing and transitions throughout the task, relying on the memory of spatial locations learned during the earlier parts.

#### No vision and eyes-closed

Participants kept their eyes closed but continued to move them as in the previous parts, following the same fixation patterns. The three-tone sequence guided their eye movements and timing.

### Anatomical data preprocessing

Results included in this manuscript are derived from anatomical preprocessing performed using *fMRIPrep* version 23.1.4 (Esteban et al., 2019) which is built on *Nipype* 1.8.6 (Gorgolewski et al., 2011). Each participant’s T1-weighted (T1w) structural image was corrected for intensity non-uniformity (INU) using N4BiasFieldCorrection (Tustison et al., 2010), implemented via ANTs (Avants et al., 2008), and was used as the T1w-reference in the workflow. The T1w-reference was skull-stripped using the *antsBrainExtraction.sh* pipeline from ANTs, with the OASIS30ANTs template as the target. Brain tissue segmentation into cerebrospinal fluid (CSF), white matter (WM), and gray matter (GM) was performed on the brain-extracted T1w using *FAST* algorithm (Zhang et al., 2001). Brain surfaces were reconstructed using FreeSurfer’s *recon-all* (version 7.3.2;(Dale et al., 1999), and the brain mask estimated previously was refined with a custom variation of the method to reconcile ANTs-derived and FreeSurfer-derived segmentations of the cortical gray-matter of Mindboggle (Klein et al., 2017). T1-weighted structural images were defaced using *PyDeface* (github.com/poldracklab/pydeface).

### Functional data preprocessing

Functional data preprocessing was performed using *fMRIPrep* 23.1.4 (Esteban et al., 2019), which is based on *Nipype* 1.8.6 (Gorgolewski et al., 2011). First, a reference volume and its skull-stripped version were generated by aligning and averaging 1 single-band references (SBRefs). Head-motion parameters with respect to the BOLD reference (transformation matrices, and six corresponding rotation and translation parameters) are estimated before any spatiotemporal (Jenkinson et al., 2002). The estimated *fieldmap* was then aligned with rigid-registration to the target EPI (echo-planar imaging) reference run. The field coefficients were mapped on to the reference EPI using the transform. BOLD runs were slice-time corrected to 0.56 s (0.5 of slice acquisition range 0 s – 1.12 s) using 3dTshift from AFNI (Cox & Hyde, 1997). The BOLD reference was then co-registered to the T1w reference (FreeSurfer) which implements boundary-based registration (Greve & Fischl, 2009). Co-registration was configured with twelve degrees of freedom to account for distortions remaining in the BOLD reference.

### Eye-tracking data preprocessing

Eye-tracking data was first converted using eye2bids (github.com/bids-standard/eye2bids) to comply with BIDS standards (Szinte et al., 2026). Blinks were removed and interpolated by excluding samples 0.15 s before and after periods when pupil size was null. The raw gaze data was converted to degrees of visual angle and centered at the screen midpoint. Slow signal drift was removed using linear detrending during periods when participants were fixating on the screen center. Finally, the eye-tracking signal was downsampled to 1 point per TR (1.2 s) in order to compare it to output from DeepMReye (see below). Note that this single-point representation was used for evaluation only.

### Gaze decoding

DeepMReye (Frey et al., 2021) is a convolutional neural network that decodes gaze position directly from MR signals of the eyeballs during fMRI scanning (Figure 1B). DeepMReye makes eye-tracking freely available to researchers by providing their model training weights online (*dataset_1to5:* https://osf.io/mrhk9) and codes to decode gaze position and fine-tune their models (github.com/DeepMReye). We aimed to evaluate the feasibility of fine-tuning the DeepMReye model using data from our tasks, including the visuomotor calibration tasks and the visual/eyes state tasks (see Figure 1B), separately and together. To do so, for each task, we generated gaze position labels from camera-based eye-tracking data and provided them alongside collected BOLD signals for fine-tuning (see *model fine-tuning*). We then evaluated the model’s predicted gaze position against these camera-based labels (see *model evaluations*). The fine-tuned model weights for all tasks are publicly available at (https://figshare.com/s/fe874ffbb37f5bc08645).

#### Visuomotor calibration tasks: model fine-tuning

DeepMReye’s architecture requires 10 gaze positions per TR as training labels, which has been shown to improve model performance compared to training on a single gaze position per TR (Frey et al., 2021). These 10-point training labels are distinct from the single-point-per-TR labels used for evaluation (see Eye-tracking data preprocessing). In order to generate these 10 labels, we implemented a quality control procedure to ensure that only reliable data were used for model training. Beginning with raw eye-tracking data, we replaced blink periods (as defined above) with NaN values, converted coordinates to degrees of visual angle, and centered them at the screen midpoint. For each non-overlapping TR window (1.2 s), we retained only those windows where at least 50% of samples contained valid data (non-NaN values). Valid windows were down sampled to 10 coordinates per TR through averaging. Windows that failed to meet the 50% validity threshold were set to NaN and consequently excluded from the training labels. The model was fine-tuned using the following parameters: learning rate = 0.000002, batch size = 15 participants, steps per epoch = 5000, epochs = 1, validation steps = 1, and learning rate decay = 0.03 similar to earlier reports (Nau et al., 2025). Our dataset was split into five equally sized partitions containing different participants. We fine-tuned the model on four partitions and tested on the fifth partition (80%/20% train-test split). This procedure was cross-validated until all data partitions and hence all participants were tested once. All other parameters were consistent with those used in the original DeepMReye training protocol (Frey et al., 2021)

To isolate the effect of fine-tuning, we compared the predictions of: i) “DeepMReye”: Decoding using the pre-trained model only (Frey et al., 2021), ii) “*DeepMReye & visuomotor calibration”:* the fine-tuned model trained on visuomotor calibration tasks data, and ii) “*Scaled DeepMReye”:* a scaled version of the decoding using pre-trained DeepMReye model (Frey et al., 2021). Scaled DeepMReye tests whether linear scaling can match or exceed fine-tuning performance, given that the pre-trained model was trained on screens smaller than the one used in our experiments (training datasets ranging from 8 dva to 15 dva sides). We applied a leave-one-participant-out cross-validation procedure as follows. For each iteration, one participant was held out as the test set. For the remaining participants, we obtained gaze predictions from the pre-trained DeepMReye model and compared them to preprocessed eye-tracking data (see Eye-tracking data preprocessing). These participants were split between 80% training and 20% validation to match the data partitioning used in DeepMReye fine-tuning, though for this linear regression we used only the training set. We fit two separate linear regression models (one for the x dimension and one for the y dimension) to map the pre-trained model’s predictions to actual gaze positions. These scaling parameters were then applied to the held-out test participant’s predictions. This procedure was repeated for each participant, ensuring that scaling corrections were derived independently from test data.

#### Visuomotor calibration tasks: model evaluations

To evaluate decoding performance, we computed two metrics for each of the three models. First, we calculated the 2D Euclidean error between predicted gaze positions, obtained as the median across the 10 predictions per TR, and preprocessed eye-tracking data at each timepoint. Second, we computed Pearson correlations separately for the x and y dimensions between predicted gaze positions and preprocessed eye-tracking data, then averaged these two correlations to obtain a final correlation score for each model. We compared three models: “*DeepMReye*”, “*DeepMReye & visuomotor calibration*”, and “*Scaled DeepMReye”*.

#### Visual/eyes state tasks: model fine-tuning

The “*DeepMReye & visual/eyes state*” model was trained starting from the original pre-trained weights using the same parameters and cross-validation procedure as described for the visuomotor calibration tasks fine-tuning. Training labels were generated following the quality control procedure described above, with specific handling for each part. For the “Vision and eyes-open” and “No vision and eyes-open” parts, labels were provided for all TR windows meeting the 50% validity threshold. For the “Vision and eyes-blink” part, labels during the prolonged blink periods (between the second and third tones) were set to NaN, meaning the corresponding fMRI data were included in training but no gaze position labels were provided for these timepoints. For the “No vision and eyes-closed” part, all labels were set to NaN throughout the entire part, as eye-tracking data were unavailable when eyes remained closed. To test whether these unlabeled eyes-closed epochs contributed to the observed fine-tuning benefit, we trained an additional control model, “*DeepMReye & visual/eyes state, target labels*”, using the same architecture, parameters, and cross-validation procedure as above, but with proxy target-position labels assigned to the eyes-closed epochs based on the known coordinates of the fixation targets, rather than leaving them unlabeled.

#### Visual/eyes state tasks: model evaluations

Since ground-truth gaze coordinates are unavailable when eyes are closed, we assessed whether decoded gaze features preserved the spatial structure of the tasks using a classification approach. Specifically, we trained a logistic regression classifier with L2 regularization to predict triangle rotation direction (one of four orientations) from decoded gaze patterns. Each decoded gaze feature comprised 18 gaze coordinates (x₁–x₉, y₁–y₉) corresponding to the three positions per triangle that were each fixated for three TRs. We evaluated classification performance using both the original temporally-ordered gaze coordinates and a temporally-shuffled version, where coordinate order was randomized. The temporally-shuffled condition removed temporal sequence information to isolate spatial shape-based decoding, while the temporally-ordered condition preserved temporal sequence information. Classification was performed separately for each experimental condition using leave-one-participant-out cross-validation, where each participant was held out once for testing, while the remaining participants were used for training. Model performance was evaluated using classification accuracy and confusion matrices averaged across participants for all four models: *’DeepMReye’, ‘DeepMReye & visuomotor calibration*’, *’DeepMReye & visual/eyes state’*, and ‘*DeepMReye & visual/eyes state, target labels*’.

### Eyelid-state decoding

#### Eyelid-state decoding: model fine-tuning

We finally evaluated the ability of DeepMReye to classify the state of the eyelids (open versus closed) across different fine-tuning strategies. The model was configured to predict 1D continuous output (with a dummy second dimension retained for architectural compatibility) rather than 2D gaze coordinates. Training labels were generated by calculating the proportion of time the eyelids were closed within each TR window using camera-based eye-tracking data. Blinks were identified as periods where pupil size equaled zero, and the proportion of samples with zero pupil size was computed for each non-overlapping TR window, yielding continuous labels ranging from 0 (eyes fully open) to 100% (eyes fully closed). We compared the same three model configurations as described above: “*DeepMReye*” (weights: *dataset_6:* https://osf.io/mrhk9), “*DeepMReye & visuomotor calibration*” (weights from our visuomotor calibration tasks), and “*DeepMReye & visual/eyes state*” (weights from our visual/eyes state tasks). Fine-tuning was performed using the same parameters and cross-validation procedure as in the gaze decoding implementation, except for Euclidean error loss weight of 1 and confidence loss weight of 0, as the model predicted continuous eye closure proportions rather than spatial coordinates.

#### Eyelid-state decoding: model evaluations

To evaluate model performance, we first assessed whether the models could reliably recover the proportion of time spent with eyelids-closed from each TR volume. First, the continuous prediction of DeepMReye were compared to ground-truth labels using Pearson correlation per run and per participant, averaged over runs. To assess discrete classification performance, we then converted the continuous predicted timeseries into binary labels by applying a 30% threshold: TRs for which the predicted eyelids-closed percentage exceeded 30% were assigned the label “eyelids-closed,” while those at or below 30% were assigned the label “eyelids-open.” The same threshold was applied to the ground-truth pupil-size-derived signal to generate the corresponding binary reference labels. Classification accuracy was then computed by comparing these two binarized timeseries.

## Statistical analysis

Statistical significance was assessed using non-parametric permutation tests across participants. For each comparison, we randomly shuffled participant labels 10,000 times to generate a null distribution of the test statistic under the assumption that participants could be interchanged without affecting the outcome. (one-tailed, testing whether fine-tuned models showed reduced error or improved performance relative to baseline models). To account for multiple comparisons, we defined pre-specified hypothesis families for the “Visuomotor calibration tasks”, the “’Visual/eyes state tasks” as well as the “Eyelid-state analysis” and applied Benjamini-Hochberg false discovery rate (FDR) correction within each (α = 0.05). Each set of tests consists of one comparison between fine-tuned and baseline models, per experiment, per measurement.

## Results

The results are organized around two objectives. First, we evaluate whether fine-tuning DeepMReye on visuomotor calibration data improves gaze decoding when the eyes are open (“Visuomotor calibration tasks”). Second, we assess whether gaze position and eyelid-state can be decoded during no-vision and eyes-closed parts, using a novel auditory-guided paradigm (“Visual/eyes state tasks”). Both tasks are illustrated in Figure 1.

### Fine-tuning improves DeepMReye performance when the eyes are open

To evaluate whether DeepMReye’s decoding performance for eyes-open parts could be adjusted to the experimental setup, we used a visuomotor calibration tasks composed of three parts: guided fixation, guided pursuit, and free viewing (Figure 1A). We then compared two model configurations: “*DeepMReye*” (the original pre-trained model, see Figure 2A) and “*DeepMReye & visuomotor calibratio*n” (fine-tuned on the visuomotor calibration tasks, see Figure 2B). Decoding accuracy was quantified in comparison to camera-based eye-tracking data using both Pearson correlation, tracking temporal dynamics, and Euclidean errors, quantifying the spatial distance between decoded and eye-tracking gaze position.

**Figure 2.**
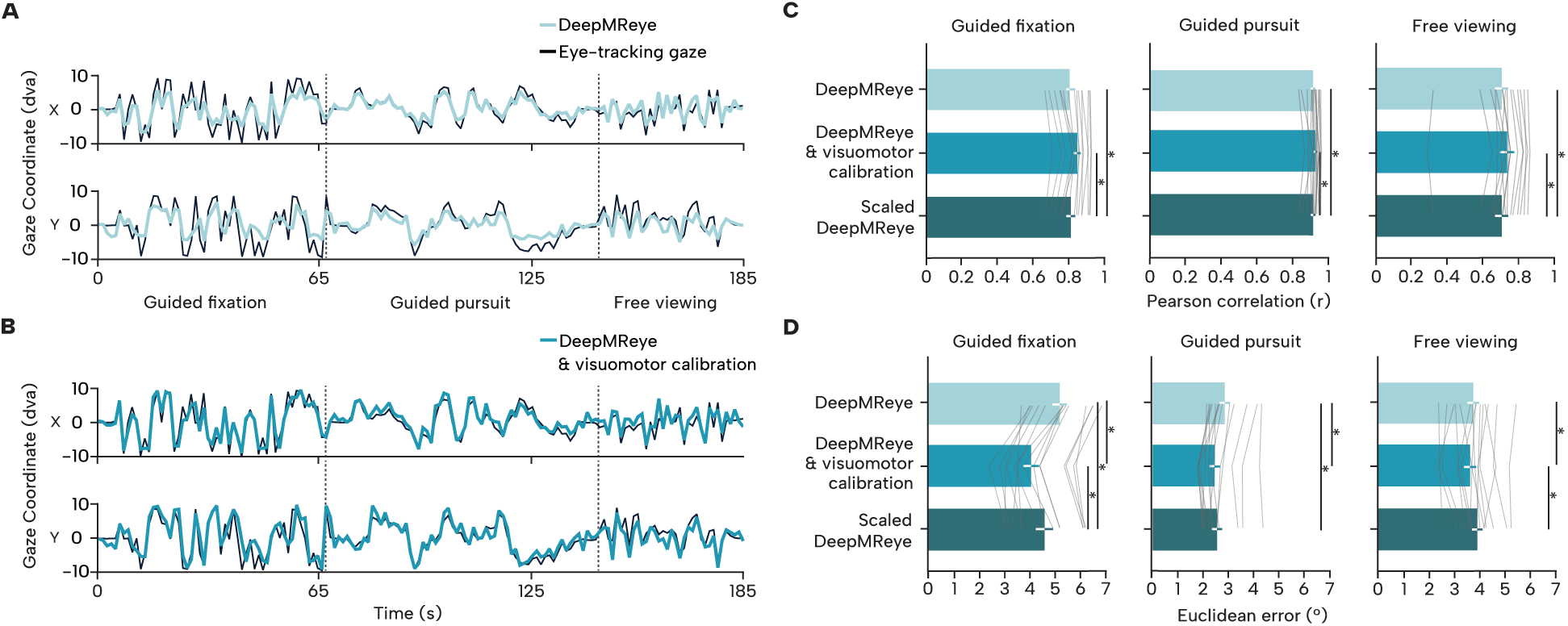
Visuomotor calibration tasks results. **A-B**. Example timeseries of the visuomotor calibration tasks of gaze coordinates decoded by *DeepMReye* (A) and *DeepMReye & visuomotor calibration* models (B), compared to downsampled eye-tracking measures. Upper and lower panels show respectively horizontal (X) and vertical (Y) coordinates of a entire run for a representative participant (sub-03). **C-D.** Pearson correlation (C) and Euclidian error (D) between decoded and eye-tracking gaze position for *DeepMReye*, *DeepMReye & visuomotor calibration*, and *Scaled DeepMReye* models for the guided fixation (left panels), guided pursuit (center panels), and free viewing parts (right panels). Bar shows mean across participants, error bar shows bootstrapped 95% CIs across participants, asterisk shows comparisons significant after Benjamini-Hochberg FDR correction within the hypothesis.

When considering temporal Pearson correlation *“DeepMReye & visuomotor calibration*” significantly outperformed the pretrained version of *“DeepMReye”* in the fixation part (Figure 2C, “*DeepMReye & visuomotor calibration*”: r = 0.87, [0.84; 0.91]; – mean Pearson and bootstrapped 95 confidence interval across participants – vs. “*DeepMReye*”: r = 0.84, [0.79; 0.88]; *p < 0.05*), the guided pursuit part (*“DeepMReye & visuomotor calibration”*: r = 0.93, [0.91; 0.94]; vs. *“DeepMReye”*: r = 0.91, [0.90; 0.93]; *p < 0.05*) and the free viewing part (*“DeepMReye & visuomotor calibration”*: r = 0.73, [0.65; 0.79]; vs. *“DeepMReye”*: r = 0.70, [0.63; 0.75]; *p < 0.05*). Similar improvements were observed when considering Euclidian errors. *“DeepMReye & visuomotor calibration*” outperformed the pretrained-only *“DeepMReye”* in the guided fixation part (Figure 2D, “*DeepMReye & visuomotor calibration*”: 4.04°, [3.53; 4.60]; –mean Euclidian errors and bootstrapped 95 confidence interval across participants – vs. “*DeepMReye*”: 5.18°, [4.69; 5.68]; *p < 0.05*), the guided pursuit part (*“DeepMReye & visuomotor calibration”*: 2.43°, [2.12; 2.81]; vs. *“DeepMReye”*: 2.84°, [2.51; 3.21]; *p < 0.05*) but not in the free viewing part (*“DeepMReye & visuomotor calibration”*: 3.63°, [3.24; 4.03]; vs. *“DeepMReye”:* 3.76°, [3.38; 4.12]; *p = 0.155*). These results show that fine-tuning DeepMReye on dedicated visuomotor calibration data improves eyes-open decoding performance, after correction for multiple comparisons, on experimental setups that were not used in the original training data.

### Fine-tuning extends the effective range of DeepMReye beyond its original training data

Having established that fine-tuning improves decoding accuracy, we next examined what drives these improvements. Because DeepMReye was originally trained on data restricted to 10 degrees of visual angle (dva) square, while our apparatus used an 18 dva square, fine-tuning may primarily help by compensating for this range mismatch. To test this, we compared model performance using only decoded gaze data within a 10 dva square. When considering Pearson correlation *“DeepMReye & visuomotor calibration*” performed slightly better than *“DeepMReye”* in the guided fixation part (*“DeepMReye & visuomotor calibration”*: r = 0.75, [0.68; 0.82]; vs. *“DeepMReye”*: r = 0.69, [0.61; 0.77]; *p* < 0.05), while no differences were observed in the guided pursuit part (*“DeepMReye & visuomotor calibration”*: r = 0.88, [0.85; 0.89]; vs. *“DeepMReye”*: r = 0.85, [0.81; 0.89]; *p* = 0.129) and in the free viewing part (*“DeepMReye & visuomotor calibration”*: r = 0.61, [0.53; 0.67]; vs. *“DeepMReye”*: r = 0.59, [0.53; 0.64]; *p* = 0.104). Opposite results were found when considering Euclidian error, with “*DeepMReye & visuomotor calibration*” performing worse than *“DeepMReye”* in the guided fixation part (*“DeepMReye & visuomotor calibration”*: 3.42°, [2.91; 3.96]; vs. *“DeepMReye”*: 3.03°, [2.60; 3.47]; *p* < 0.05) while the guided pursuit part did not reach significance after correction (*“DeepMReye & visuomotor calibration”*: 2.11°, [1.91; 2.36]; vs. *“DeepMReye”*: 1.99°, [1.79; 2.23]; *p* = 0.055), and no difference were observed in the free viewing part (*“DeepMReye & visuomotor calibration”*: 3.26°, [2.92; 3.53]; vs. *“DeepMReye”*: 3.09°, [2.85; 3.31]; *p* = 0.0891). These findings indicate that for decoding gaze within 10 dva square range, the pre-trained DeepMReye model is more accurate in line with its training-data range. Consequently, fine-tuning is most beneficial for decoding gaze positions that lie outside the range of the original training data.

### Fine-tuning effectiveness extends beyond scaling adjustments

One possible explanation for these range-dependent improvements is that fine-tuning simply learns to rescale the model output to match the larger screen dimensions of our apparatus. To test this, we compared the fine-tuned model against a scaled version of DeepMReye that explicitly accounted for our larger screen dimensions (see Material and methods). This analysis evaluated whether visuomotor calibration offered advantages beyond linear scaling of the gaze decoding output. When considering temporal Pearson correlation, “*DeepMReye & visuomotor calibration*” performed better than “Scaled DeepMReye” in the guided fixation part (Figure 2C*, “DeepMReye & visuomotor calibration”*: r = 0.87, [0.84; 0.91]; vs. *“Scaled DeepMReye”*: 0.84, [0.79; 0.88]; *p* < 0.05), the guided pursuit part (*“DeepMReye & visuomotor calibration”*: r = 0.93, [0.91; 0.94]; vs. *“Scaled DeepMReye”*: 0.91, [0.90; 0.93]; *p* < 0.05), as well as the free viewing part (*“DeepMReye & visuomotor calibration”*: r = 0.73, [0.65; 0.79]; vs. *“Scaled DeepMReye”*: r = 0.70, [0.63; 0.75]; *p* < 0.05). While considering Euclidean error, “*DeepMReye & visuomotor calibration*” performed better than “Scaled DeepMReye”, in the guided fixation part (Figure 2D, *“DeepMReye & visuomotor calibration”*: 4.04°, [3.53; 4.60]; vs. *“Scaled DeepMReye”*: 4.57°, [4.00; 5.19]; *p* < 0.05) and the free viewing part (*“DeepMReye & visuomotor calibration”*: 3.26°, [2.92, 3.53]; vs. *“Scaled DeepMReye”*: 3.92°, [3.56; 4.28]; *p* < 0.05), while no difference was found in the guided pursuit part (*“DeepMReye & visuomotor calibration”*: 2.43°, [2.12; 2.81]; vs. *“Scaled DeepMReye”*: 2.53°, [2.24; 2.89]; *p* = 0.088). These results suggest that the beneficial effects of fine-tuning on model performance go beyond the learning of output scaling and include a better tracking of the dynamics in gaze behavior.

### Camera-based eye-tracking data are not required for model training

Finally, we asked whether fine-tuning requires access to camera-based eye-tracking data, a resource that may not be available in all settings. To address this, we compared model performance when trained using target position labels instead of eye-tracking labels. This was done for the guided fixation and guided pursuit part, which were the only parts of the visuomotor calibration tasks in which labels could be provided during the training process. First, considering temporal Pearson correlations, differences between the training schemes were generally small. The model trained on target position showed slightly lower correlation only in the guided fixation part (“*Target position labels*”: r = 0.88, [0.84; 0.91]; vs. “*Eye-tracking labels*”: r = 0.87, [0.84; 0.91]; *p* < 0.05). No significant differences were observed in the guided pursuit part (*p* = 0.419). Similarly, when considering Euclidean error, decoding performance was comparable between models trained on target position labels and eye-tracking labels, with only a modest reduction observed for the guided fixation part (“*Target position labels*”: 3.89°, [3.35; 4.45]; vs. “*Eye-tracking labels*”: 4.04°, [3.53; 4.60]; *p* < 0.05), while no difference was observed in the guided pursuit part (“*Target position labels*”: 2.48°, [2.18; 2.84]; vs. “*Eye-tracking labels*”: 2.43°, [2.12; 2.81]; *p* = 0.089). These results demonstrate that DeepMReye can be fine-tuned using target-position labels without substantial differences in decoding accuracy, highlighting its potential for use in settings where eye-tracking is not available.

Taken together, the results of the visuomotor calibration tasks demonstrate that DeepMReye can be effectively fine-tuned with a small participant sample (N = 15) to a specific experimental apparatus, yielding substantially improved decoding performance, that extends the decoding range without simply scaling the output. Crucially, this fine-tuning does not require camera-based eye-tracking data, making it accessible to all laboratories. Having established robust gaze decoding under eyes-open parts, we next ask whether similar decoding is possible when no visual input is available and the eyes are closed.

### Decoding gaze when the eyes are closed

Having demonstrated that DeepMReye can be fine-tuned using eyes-open visuomotor calibration tasks, we investigated decoding gaze position when the eyes are closed. To this end, we developed a set of novel fMRI tasks, the visual/eyes state tasks, in which participants performed guided eye movements tracing one of four triangle shapes, each oriented in a different cardinal direction (up, down, left, right). Each corner of the triangle was paired with a distinct auditory cue, allowing participants to associate temporal auditory information with spatial positions (Figure 1C).

The visual/eyes state tasks comprised of four parts that systematically varied the visual input (vision, no vision) and eyes-state (eyes-open, eyes-blink, eyes-closed): (1) “*Vision with eyes-open*”, in which participants viewed a screen and followed auditory cues with open eyes, (2) “Vision and eyes-blink”, where participants blinked during each fixation point guided by auditory cues, (3) “No vision and eyes-open”, in which no stimulus was shown but the eyes remain open, and lastly (4) “No vision and eyes-closed”, in which participants replicated the same sequences with their eyes closed, guided solely by auditory cues. Importantly, this design enabled the systematic evaluation of DeepMReye’s decoding performance during eyes-closed states using structured, auditory-guided eye movements, going beyond the single-participant proof-of-concept reported previously (Frey et al., 2021; Nau et al., 2025). Moreover, it allowed us to assess the decoding performances further in all parts by determining the accuracy of a logistic-regression classifier in decoding the direction of the triangle obtained by the decoded eye traces.

### No fine-tuning improvements when considering temporally-ordered coordinates

We evaluated decoding performance by training a logistic-regression classifier on the gaze traces decoded by each model configuration and assessing its accuracy in identifying the direction of the triangle. As a first step, we used the temporally-ordered coordinate sequences, that is, the decoded gaze positions in the exact order in which they were sampled during the task. Because the triangle shapes were traced in a fixed sequence, a classifier can in principle exploit the temporal order of the coordinates alone to identify the triangle direction, regardless of the spatial accuracy of the decoded gaze positions. Under this condition, classification accuracy was high across all model configurations, with no significant differences between models for the “Vision and eyes-open” part (“*DeepMReye*”: 94.72%, [84.17; 100.00]; –mean accuracy and bootstrapped 95 confidence interval across participants– vs. “*DeepMReye & visuomotor calibration*”: 100%, [100.00; 100.00]; vs. (“*DeepMReye & visual/eyes state*”: 100%, [100.00; 100.00]; *p = 0.5012*) and the “Vision while blinking” part (“*DeepMReye*”: 95.83%, [87.50; 100.00]; vs. “*DeepMReye & visuomotor calibration*”: 100%, [100.00; 100.00]; vs. “*DeepMReye & visual/eyes state*”: 100%, [100.00; 100.00]; *p = 0.5012*). In the “No vision and eyes-open” part, classifier accuracy was slightly lower overall yet it showed negligible differences between models (“*DeepMReye*”: 88.06%, [75.56; 97.78]; vs. “*DeepMReye & visuomotor calibration*”: 88.89%, [77.78; 97.22]; vs. “*DeepMReye & visual/eyes state*”: 90.56%, [81.11; 97.78]; *p = 0.5012*). The same pattern was observed in the “No vision and eyes-closed” part (“*DeepMReye*”: 85.83%, [76.39; 93.61]; vs. “*DeepMReye & visuomotor calibration*”: 86.11%, [77.22; 93.06]; vs. “*DeepMReye & visual/eyes state*”: 86.11%, [78.05; 93.33]; *p = 0.5013*). While encouraging, these ceiling-level accuracies likely reflect the exploitation of temporal structure rather than true gaze decoding performance, and therefore do not provide a sensitive test of fine-tuning improvements. (Figure 3A).

**Figure 3.**
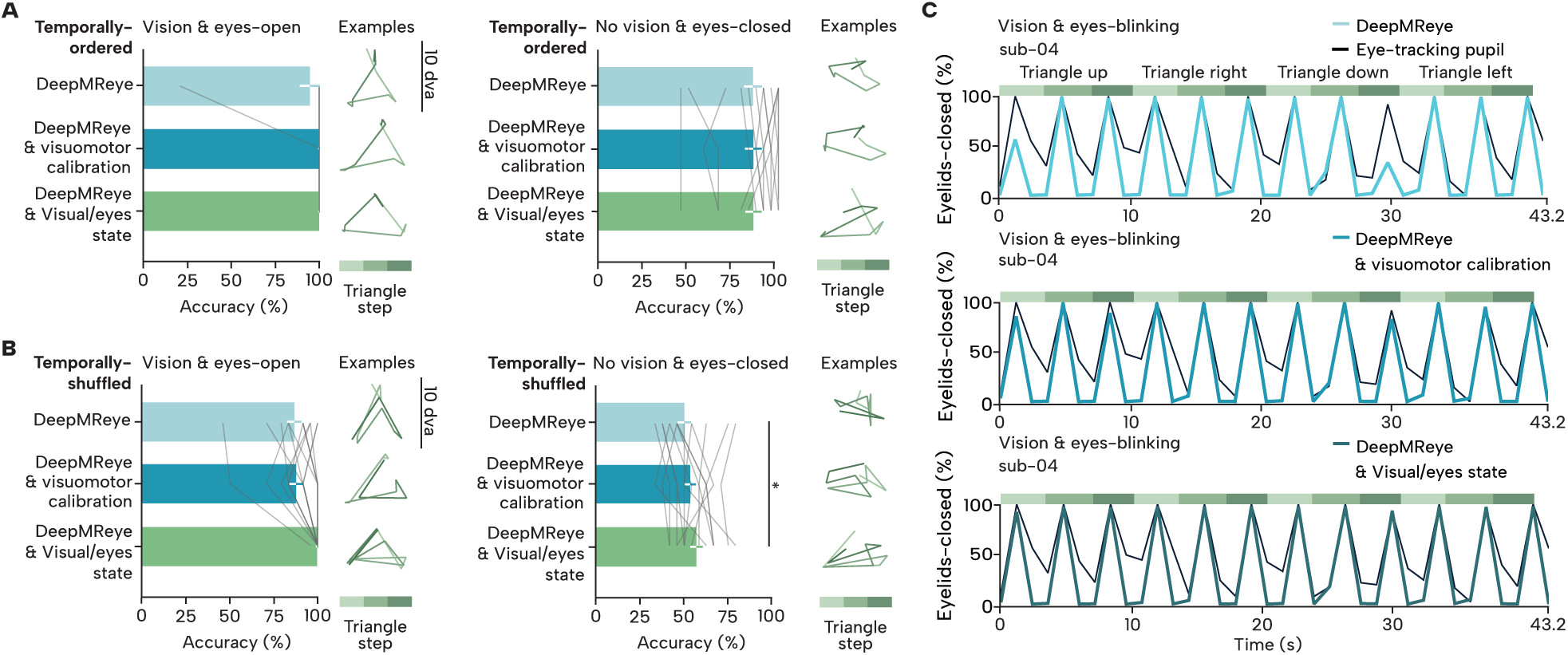
**A.** Classification accuracy for “*DeepMReye”*, “*DeepMReye & visuomotor calibration”*, and “*DeepMReye & visual/eyes state”* models using temporally-ordered gaze coordinates, shown for the “Vision and eyes-open” and “No vision and eyes-closed” parts. Example segments (triangle-up trial from sub-04) illustrate the temporally-ordered or temporally-shuffled gaze coordinate inputs used by each model configuration. **B.** Classification accuracy for the same model configurations and parts following temporally-shuffled coordinates. Error bars represent bootstrapped 95% confidence intervals across participants; asterisk shows comparisons significant after Benjamini-Hochberg FDR correction within each hypothesis. **C.** Timeseries of one triangle sequence of a representative participant (sub-04) of the proportion of eyelid-closed per sample during “Vision and eyes-blink” part, as decoded by “*DeepMReye*” (top panel), *“DeepMReye & visuomotor calibration”* (middle pannel), and “*DeepMReye & visual/eyes state*” (bottom panel), compared to the ground-truth training labels.

### Temporally-shuffled coordinates reveal fine-tuning effects on eyes-closed data

As we reasoned that the above results reflect more the temporal structure of the experiment than the actual decoding performance, we next shuffled the temporal order of the decoded gaze coordinates before classification. By removing the sequential structure of the eye-movement traces, the classifier can no longer rely on temporal order and must instead decode triangle direction from the spatial distribution of gaze positions alone. In the “Vision and eyes-open” part, fine-tuning substantially improved classification accuracy (“*DeepMReye*” vs. “*DeepMReye & visual/eyes state*”: 99.72%, [99.17; 100.00]; *p < 0.05*), while calibration-only fine-tuning did not reach significance after correction (“*DeepMReye & visuomotor calibration*”: 87.78%, [80.00; 93.61]; *p = 0.497*). Similarly, in the “No Vision and eyes-closed” part, fine-tuning with visual/eyes state data significantly improved performance (“*DeepMReye*” vs. “*DeepMReye & visual/eyes state*”: 56.94%, [51.39; 63.06]; *p < 0.05*), while calibration-only fine-tuning again did not reach significance (’DeepMReye & visuomotor calibration’: 53.33%, [47.78; 58.62]; *p = 0.257*). For the “Vision and eyes-blink” part, fine-tuning with both visuomotor calibration data and visual/eyes state data improved classification accuracy compared to the pre-trained model (“*DeepMReye*”: 80.83%, [72.22; 88.06]; vs. “*DeepMReye & visuomotor calibration*”: 90.83%, [87.50; 93.89]; *p < 0.05*; “*DeepMReye*” vs. “*DeepMReye & visual/eyes state*”: 94.72% [91.94; 97.50]; *p < 0.05)*). Task-specific fine-tuning improved gaze decoding accuracy after the gaze coordinates were shuffled, after correction. Calibration-only fine-tuning improved performance only in the eyes-blink part.

### Labelling eyes-closed epochs provides no additional decoding benefit

Finally in order to directly test whether including proxy labels, derived from the known target position, for the eyes-closed parts improves the observed fine-tuning effects, we compared two fine-tuned models: one trained with the eyes-closed epochs left unlabeled (“DeepMReye & visual/eyes state”), and one trained with proxy target-position labels assigned to the eyes-closed epochs (“DeepMReye & visual/eyes state, target labels”). Using the classification approach described above, we found no significant difference in decoding accuracy between the two models in the temporally-shuffled condition of the “No vision and eyes-closed” part (“DeepMReye & visual/eyes state”: 56.94%, [51.39; 63.06]; vs. “DeepMReye & visual/eyes state, target labels”: 53.21% [48.08; 58.97]; *p = 0.966*). This null result held across all four task parts and both temporally-ordered and temporally-shuffled conditions (all p ≥ 0.75). One potential explanation for why including eyes-closed training data did not improve model performance is that proxy labels for eyes-closed epochs rely on an unverifiable assumption of compliance. Potentially, eyes-closed epochs may simply add noise rather than improve the model further.

### Eyelid-state decoding

Beyond gaze position, we asked whether fine-tuning also improves decoding of eyelid-state, specifically whether the model can accurately distinguish between open and closed eyelids. Importantly, this analysis also addresses a more practical question: does eyelid-state decoding require task-specific fine-tuning with eyes-closed data, or is fine-tuning with eyes-open visuomotor calibration data sufficient?

To evaluate this, we compared three model configurations using eye-tracking pupil size as ground-truth. Because other parts of the visual/eyes state tasks were heavily imbalanced toward either open or closed eyelids periods, we focused our analysis on the “Vision and eyes-blink” part, which provided the most balanced distribution of eyelid-state periods and therefore the most sensitive test of decoding performance.

Classification accuracy was computed by converting both the continuous model predictions and the eye-tracking ground truth into binary labels using a 30% threshold: any sample with an eye-closure percentage above 30% was assigned the label “eyelid-closed,” while any sample at or below 30% was assigned the label “eyelids-open.” This binarization was applied identically to both the predicted and ground-truth signals before computing accuracy. Figure 3C show an example timeseries of one triangle sequence of a representative participant (sub-04) of the proportion of eyelids-closed per sample during “Vision and eyes-blink” part of the visual/eye state tasks. In the “Vision and eyes-blink” part, fine-tuning improved eyelid-state decoding accuracy compared to the pre-trained model (“*DeepMReye*”: 76.22%, [72.41; 79.32]; vs. “*DeepMReye & visuomotor calibration*”: 78.86%, [74.92; 82.52]; *p* < 0.05; “*DeepMReye*” vs. “*DeepMReye & visual/eyes state*”: 79.78%, [76.71; 82.58]; *p* < 0.05). A similar pattern was observed for correlation with the eye-tracking pupil measures, yet after correction there was no significance found (“*DeepMReye*”: r = 0.86, [0.74; 0.94]; vs. “*DeepMReye & visual/eyes state*”: r = 0.87, [0.77; 0.9]; p *= 0.064*). Critically, “*DeepMReye & visuomotor calibration*” and “*DeepMReye & visual/eyes state*” did not significantly differ on any metric (all *p* > 0.17), with mean accuracy differences of less than 1%. These results indicate that fine-tuning on eyes-open visuomotor calibration data alone is sufficient for eyelid-state decoding, and that additional fine-tuning with eyes-closed data provides no additional benefit.

## Discussion

Eye-tracking in an MRI scanner is technically demanding, and many laboratories lack the equipment to do it at all. Here, we asked whether a short visuomotor calibration paradigm with a set of classical oculomotor tasks can be used to fine-tune DeepMReye and adapt it to an apparatus that was not part of the original data used for pretraining. Indeed, we found that fine-tuning reliably improved decoding across a wider-than-standard visual field (18 dva-square), and that this improvement reflected genuine model adaptation rather than simple output rescaling. Going beyond the abilities of camera-based eye-tracking, we then examined eyes-closed conditions. Notably, we show that complex, structured eye movements performed with closed eyes can be decoded from fMRI data. Using an auditory-guided paradigm, we generated eyes-closed data for the model to predict and showed that task-specific fine-tuning substantially improved decoding performance. This opens a window into paradigms where eyes must remain closed, such as sleep, resting-state, and visual imagery tasks, as well as tasks performed with populations for whom camera-based eye-tracking has never been a realistic option (patient, children). Finally, we also show that DeepMReye can differentiate open and closed eyelids with fine-tuned models outperforming the pre-trained baseline.

## Practical Considerations for Researchers

Several findings from this study have direct practical implications for researchers wishing to deploy DeepMReye in their own apparatus.

### When to fine-tune, and when not to

Fine-tuning consistently improved decoding specifically when gaze was expected to extend into the periphery and beyond the range of the original training data. However, surprisingly, it worsened performance when analyses were restricted to the central 10 degrees of visual angle square, the range on which DeepMReye was originally trained. Researchers whose paradigms keep gaze within this central region may therefore be advised to use the pre-trained DeepMReye model without any fine-tuning, while those using peripheral targets or wide-field stimuli should invest in a visuomotor calibration session. Importantly, our results indicate that linear rescaling does bring improvements, yet, it falls short of what a dedicated fine-tuning can achieve.

### Camera-based eye-tracking is not required for fine-tuning

A practical barrier in many laboratories is the lack of MR-compatible eye-tracking equipment. Our results show that using the known spatial coordinates of targets as training labels, rather than actual measured gaze coordinates, yields comparable fine-tuning performance. This approach is viable for any structured task where participants are expected to look at known locations. However, it cannot be applied to free-viewing or other naturalistic paradigms where actual gaze cannot be assumed to match target positions; for those cases, a camera-based eye tracker or Electrooculography (EOG) remain the better option.

### Eyes-closed decoding benefits from task-specific fine-tuning, not eyes-closed labels

Our results indicate that improving decoding during eye closure depends on how closely the fine-tuning data matches the task and apparatus. Generic eyes-open visuomotor calibration alone did not improve eyes-closed decoding significantly, whereas fine-tuning on the labeled eyes-open and blink portions of the same task paradigm did. This benefit was not enhanced by additionally labelling the eyes-closed epochs with proxy target positions. Researchers planning to monitor gaze behavior during eyes-closed states, for example during mental imagery, sleep, or resting-state paradigms, should therefore prioritize collecting a task– and apparatus-matched calibration run performed with the eyes open and or eyes blinking, such as the eyes-open portion of our auditory-guided paradigm, over attempting to generate labels for the eyes-closed epochs themselves. Our auditory-guided triangle-tracing task proofs useful as an evaluation tool, allowing researchers to assess eyes-closed decoding performance in the absence of camera-based ground truth. For populations or study designs in which running any structured, task-matched eyes-open calibration is difficult, the training weights shared alongside the present work provide a good starting point.

### Eyelid-state tracking has a temporal resolution ceiling

DeepMReye predicts at the sample level of a TR, making it unsuitable as a replacement for high-temporal-resolution blink detection. Its eyelid-state output is best interpreted as a coarse signal distinguishing sustained open versus closed eyelids periods, useful for identifying sleep stages, prolonged closures, or states of wakefulness in general, but not for detecting individual blinks in natural conditions.

## Further considerations for decoding eyes-closed eye-movements

The ability to decode gaze during eyes-closed periods opens a window into oculomotor behaviors that have been difficult to study non-invasively. Promising applications include sleep neuroimaging, where REM eye movements could be tracked as markers of cognitive processing (Senzai & Scanziani, 2022, Nau et al., 2025); eyes-closed resting-state and mental imagery paradigms, where eye movements reflect internally directed cognition even without visual input, and clinical populations for whom camera-based eye-tracking or sustained eyes-open fixation is not feasible. Yet several important considerations shape how these results should be interpreted.

First, it is known that eyes-closed oculomotor behavior is qualitatively different from eyes-open one. When the eyelids close, it physically deforms the eyeball, altering its shape and therefore the MR signal geometry that DeepMReye relies upon (Brodoehl et al., 2016). Beyond the mechanical effect of the eyelids, the BOLD signal itself may differ between eyes-open and eyes-closed states (Marx et al., 2004). Additionally, eyes-closed eye movements have distinct features: Bell’s phenomenon (a reflexive upward rotation of the eyes upon closure), reduced saccade frequency, and highly idiosyncratic individual movement strategies (Haladjian et al., 2015; Shaikh et al., 2010). These differences mean that a model trained primarily on eyes-open data may impose a specific structure onto decoded eyes-closed trajectories, rather than faithfully capturing the underlying movements, an open question that future work should systematically address.

At the same time, these same signal differences make eyelid-state itself decodable: the original DeepMReye model was already shown to classify eyelids open versus closed from fMRI without any fine-tuning (Frey et al., 2021), suggesting eyelid-state information is implicitly encoded in the models’ input data. Building on this, we show that fine-tuning further improves decoding, and initializing from gaze-position-fine-tuned weights is sufficient. Future work could further improve eyelid-state decoding by exploring architectures that explicitly leverage this entanglement of eyelid information in the eyeballs signal.

Second, voluntarily executing precise eye movements with eyes closed is inherently difficult (Allik et al., 1981). Participants then rely on non-visual information (auditory or proprioceptive) or on internal models of spatial locations (visuospatial memory, visual imagery) than visual cues to guide movements that are normally visually anchored (Allik et al., 1981; Balslev et al., 2011). Moreover, proprioceptive inputs are distorted due to closed eyelids and there is lack of visual feedback resulting in ocular trajectories with substantially greater spatial variability and fixation errors. Yet it has been reported, that even in darkness or with eyes closed, participants are able to report the direction of an eye movement (Skavenski et al., 1972) and produce simple geometric shapes with their eye movements (Allik et al., 1981). Still, this sets a practical ceiling on decoding accuracy that is independent of model quality. Without visual feedback, the eye movements themselves are inherently more variable: fixations are less stable, trajectories deviate more from their intended paths, and spatial errors accumulate across a sequence. A perfect decoder would faithfully recover these movements, but the recovered trajectories would still look imprecise, because the underlying movements are imprecise. As we evaluated the models’ performance implicitly through spatial accuracy, the remaining performance gap between eyes-open and eyes-closed results therefore likely reflects this greater biological variability in the movements being decoded, rather than a failure of the model itself.

## Limitation and future directions

A fundamental constraint concerns the ground-truth used to evaluate decoding performance. In eyes-open parts of our experiment, as discussed previously, we compared to real eye-tracking data as ground truth and not screen coordinates, which better captures actual gaze behavior. However, this has the implication that any noise or inaccuracy in the eye-tracker signal propagates into our performance estimates. All model performance scores are therefore a function of the accuracy of our model as well as the accuracy of the camera-based eye-tracking data itself. In eyes-closed parts, no direct ground-truth is available. We assessed decoding indirectly by training a classifier to identify the shape of the auditory-guided eye movement paradigm from decoded trajectories. While this approach provides a meaningful signal, classifier accuracy does not speak to finer-grained properties such as the precision of individual fixation positions or the fidelity of trajectory shape. The correspondence between decoded and true eyes-closed gaze trajectories remains an open empirical question. A further constraint is the visuomotor calibration task itself needed for fine-tuning for eyes-closed parts. Our auditory-guided paradigm requires participants to follow structured movement sequences reliably, which may not be feasible in all populations, including precisely those clinical populations where eyes-closed gaze decoding would be most valuable (e.g., patients with disorders of consciousness, children or specific psychiatric populations; Ting et al., 2014; Toghi et al., 2025)

These limitations suggest a practical hierarchy for laboratories working without high-quality eye-tracking data. Those with some eyes-closed recordings can still benefit from partial fine-tuning, using our released weights as a starting point. Those without any suitable calibration data could use our released weights or the original pre-trained DeepMReye model. Each level of this is preferable to having no gaze information at all.

A natural next step is improving decoding during eyes-open free viewing tasks, the condition most relevant to naturalistic experimental designs. Current performance likely reflects several compounding factors: the lack of reliable label-free training strategies, the relatively low proportion of free viewing data in the pre-trained model’s training set, and the inherent complexity of free viewing behavior itself. Crucially, many paradigms where free viewing decoding would be most valuable are precisely those where conventional eye-tracking is hardest to obtain: developmental studies with children, or naturalistic viewing in clinical populations, whose atypical gaze patterns and compliance profiles often compromise conventional eye-tracking. A camera-free MR-based approach would be particularly powerful for these groups, but will likely require retraining DeepMReye from scratch with an additional large, dedicated free viewing dataset. Alternatively, new unsupervised approaches could address this critical gap.

Moreover, the ability to decode gaze during eye closure creates new opportunities in contexts that have until now been entirely inaccessible to fMRI-based gaze tracking. Sleep neuroimaging is a particularly compelling application: continuous eye movement monitoring across sleep stages could provide a non-invasive window into REM-related oculomotor dynamics, potentially linking movement patterns cognitive processes (Senzai & Scanziani, 2022; Siclari et al., 2017). Beyond sleep, eyes-closed decoding could provide oculomotor activity during resting-state and mental imagery paradigms (Fransson et al., 2014; Ramot et al., 2011), where eye movements are known to reflect the spatial structure of internal representations even without visual input (Johansson et al., 2006; Nau et al., 2025; Wynn et al., 2019). On the clinical side, this approach is relevant for populations where conventional camera-based eye-tracking fails entirely, such as congenitally blind individuals, or yields data of insufficient quality, as is often the case in conditions associated with oculomotor disturbances such as Parkinson’s disease and multiple sclerosis (Corin et al., 1972; Serra et al., 2018). More broadly, since camera-based eye-trackers and EOG systems cannot be applied to data collected without them, the demonstration that DeepMReye can be fine-tuned without eye-tracking equipment (also see Frey et al., 2021)lowers the barrier to retrospective application of gaze decoding to archival fMRI datasets, potentially unlocking eye movement information from large existing repositories collected without simultaneous eye-tracking.

## Conclusion

Together, these results mark a meaningful step toward making fMRI-based gaze decoding broadly accessible, even for studies of eyes-closed states. By implementing practical visuomotor calibration tasks and fine-tuning procedure, and enabling structured gaze monitoring during eyes-closed states, this work expands the range of experimental contexts in which DeepMReye can serve as a practical tool. The ability to retrospectively apply these methods to existing datasets acquired without camera-based eye-tracking further underscores their wide applicability. Particularly for free-viewing paradigms and clinical populations, the advances demonstrated here lay the groundwork for a new generation of fMRI studies in which eye movements are routinely monitored regardless of whether participants’ eyes are open or closed.

## Acknowledgments

We thank the members of the INVIBE team at the Institute des Neurosciences de la Timone for valuable discussions and constructive comments on this work. We are grateful to Markus Frey for sharing code and technical expertise. We thank the MRI platform of the Institute de Neurosciences de la Timone (INT) for technical support and access to imaging facilities. We also thank all colleagues and researchers who provided helpful insights and feedback throughout the project.

## Author contributions

S.M.K. implemented and carried out the experiments, analyzed the data, and wrote the manuscript. M.N. contributed conceptual and practical advice on design and analysis, with G.S.M. contributing to their specification and refinement. M.S. supervised the project. U.L. reviewed the manuscript. All authors contributed to writing the manuscript, discussed the results, and commented on the final version.

## Ethics

This experiment was approved by the Ethics Committee of Aix-Marseille University (Comité de Protection des Personnes OUEST III, no 2018-A02608-47) in accordance with the Declaration of Helsinki. All participants gave written informed consent.

## Additional files

### Data and code availability

We make available online our imaging dataset (doi.org/10.18112/openneuro.ds006833.v1.0.0). Experimental and data analyses codes are made available online (github.com/mszinte/deepmreye). All model weights are available here (doi.org/10.6084/m9.figshare.32687379). We also provide a ready to use code repository for using the model weights here (https://github.com/sinaklg/int_deepmreye/).

